# recount: A large-scale resource of analysis-ready RNA-seq expression data

**DOI:** 10.1101/068478

**Authors:** Leonardo Collado-Torres, Abhinav Nellore, Kai Kammers, Shannon E. Ellis, Margaret A. Taub, Kasper D. Hansen, Andrew E. Jaffe, Ben Langmead, Jeffrey T. Leek

## Abstract

`recount` is a resource of processed and summarized expression data spanning nearly 60,000 human RNA-seq samples from the Sequence Read Archive (SRA). The associated `recount` Bio-conductor package provides a convenient API for querying, downloading, and analyzing the data. Each processed study consists of meta/phenotype data, the expression levels of genes and their underlying exons and splice junctions, and corresponding genomic annotation. We also provide data summarization types for quantifying novel transcribed sequence including base-resolution coverage and potentially unannotated splice junctions. We present workflows illustrating how to use recount to perform differential expression analysis including meta-analysis, annotation-free base-level analysis, and replication of smaller studies using data from larger studies. recount provides a valuable and user-friendly resource of processed RNA-seq datasets to draw additional biological insights from existing public data. The resource is available at https://jhubiostatistics.shinyapps.io/recount/.

## 1 Introduction

RNA sequencing (RNA-seq) is a ubiquitous tool for assaying gene expression. Public sequencing data repositories such as the Sequence Read Archive [1] now hold more than 50,000 human RNA-seq samples, and the size of the archive doubles approximately every 18 months (http://www.ncbi.nlm.nih.gov/Traces/sra/sra.cgi?view=announcement). Many studies in public and dbGaP-protected repositories are valuable to biological researchers and methods developers. For example, some studies are derived from individuals with rare disease [2], from hard-to-obtain tissues [3] or from rare forms of cancer [4]. Other studies are notable for their size, e.g. the GTEx study [5] consisting of 9,662 samples derived from 551 individuals and 54 body sites.

However, the majority of these archived samples are available only as compressed collections of raw sequencing reads. And while some samples have been summarized into gene counts in repositories like the Gene Expression Omnibus (GEO), these expression level summarizations are heavily dependent on the processing pipelines, which can vary dramatically across study. Typically analyses of public data require re-analysis beginning from the reads. However, processing raw reads into a form suitable for various downstream analyses is technically challenging. Care is required to craft expression summaries, derived from a common comparable pipeline - that are both concise – convenient for researchers to download and interact with – and useful in a variety of downstream scenarios.

Our first effort addressed this in two ways: (a) by summarizing data into concise gene count tables, and (b) by making processed data available in the form of Bioconductor [6] ExpressionSet objects including associated metadata using a single processing pipeline. That resource, called recount [7], was applied, for example, to development of methods for differential expression and normalization [8, 9, 10], compilation of co-expression networks [11], and to studying the effect of ribosomal DNA dosage on gene expression [12]. Here we present an updated version of recount consisting of 59,319 uniformly processed human RNA-seq samples across 2,035 projects. These publicly available SRA samples include, for example, the entirety of the GTEx, GEUVADIS [13], SEQC/MACQ-III [14], and ABRF [15] studies. Now recount summarizes the expression data across multiple feature types, including genes, exons, exon-exon splice junctions and base-level coverage. These summarizations enable a wider variety of downstream analyses, including testing for differential expression of potentially unannotated transcribed sequence. Similarly, recount now marshals relevant meta/phenotype data into a searchable interface available from both the website (https://jhubiostatistics.shinyapps.io/recount/) and the Bioconductor package https:// github.com/leekgroup/recount, allowing users to rapidly access relevant data.

We demonstrate three potential workflows using the recount database and corresponding R package. First, we show that the resource can be used to merge multiple data sets studying the same problem to perform rapid genomic meta-analyses. Next, we demonstrate that our processed version of the GTEx gene count data closely matches the official gene counts released by the project itself. This serves to demonstrate (a) that it is easy to compare the processed data from recount to data processed with other pipelines and (b) that our gene counts are consistent with those published from the GTEx project. Lastly we show that the resource can be used to easily perform differential expression analysis at different feature summarizations: exons, genes, junctions, and expressed regions [7]. We also demonstrate the ease of comparing results discovered in one study within recount to other studies in the resource for validation. All of our analyses are reproducible and the results can be compiled using the R markdown files found at: http://leekgroup.github.io/recount-analyses/.

## 2 Results

### 2.1 Data description

We analyzed the publicly available SRA and the latest (v6) release of GTEx and samples. The SRA data consisted of 49,848 publicly available samples spanning over 146 terabases of reads. These reads were downloaded and analyzed, resulting in a final set of 48,558 samples that could be fully downloaded and processed (1,290 samples could be downloaded only partially and were excluded, see Supplementary Methods). The GTEx data consisted of 9,662 samples spanning over 65 terabases of reads, 550 individuals and the K562 cell line, and 32 tissues. These reads were downloaded and analyzed, resulting in a final set of 9479 samples that could be fully downloaded and processed (183 could be downloaded only partially, see Supplementary Methods).

### 2.2 Use case 1: Meta-analysis

To illustrate the ease of combining data from multiple projects included in recount as part of a cross-study meta-analysis, we carried out a cross-tissue differential expression (DE) analysis comparing gene expression between colon and whole blood. As an initial analysis, colon samples labeled as controls were taken from studies SRP029880 (a study of colorectal cancer [16], n = 19) and SRP042228 (a study of Crohn’s disease [17], n = 41). Whole blood samples labeled as controls were taken from SRP059039 (a study of virus-caused diarrhea, unpublished, n = 24), SRP059172 (a study of blood biomarkers for brucellosis, unpublished, n = 47) and SRP062966 (a study of lupus, unpublished, n = 18). After filtering genes to include only those with an average normalized count of at least 5 across samples, we performed gene-level differential expression analysis using limma [18] and voom [9].

To validate the results, we selected all colon and whole blood samples from the GTEx project (n = 376 and 456, respectively) and performed the same analysis, adjusting for batch effects. We then computed rank-based concordance, examining the fraction of the top DE genes that were included in both analyses. Results are shown in orange in Figure 1A. Approximately 20% of the top 100 genes from the two analyses were concordant. As a comparison and to provide context for this result, we performed two additional comparisons. First, we used GTEx lung data (n = 374) in place of the colon data and computed DE genes compared to whole blood. In this case, only approximately 5% of the top 100 DE genes were shared in the top 100 genes from our multi-study analysis. Second, to represent concordance results expected for a comparison of unrelated things, we used ranked coefficients for batch instead of for tissue and see very little concordance. These comparisons support that we can use the resources found in recount to perform a valid tissue-specific meta-analysis.

**Figure 1:**
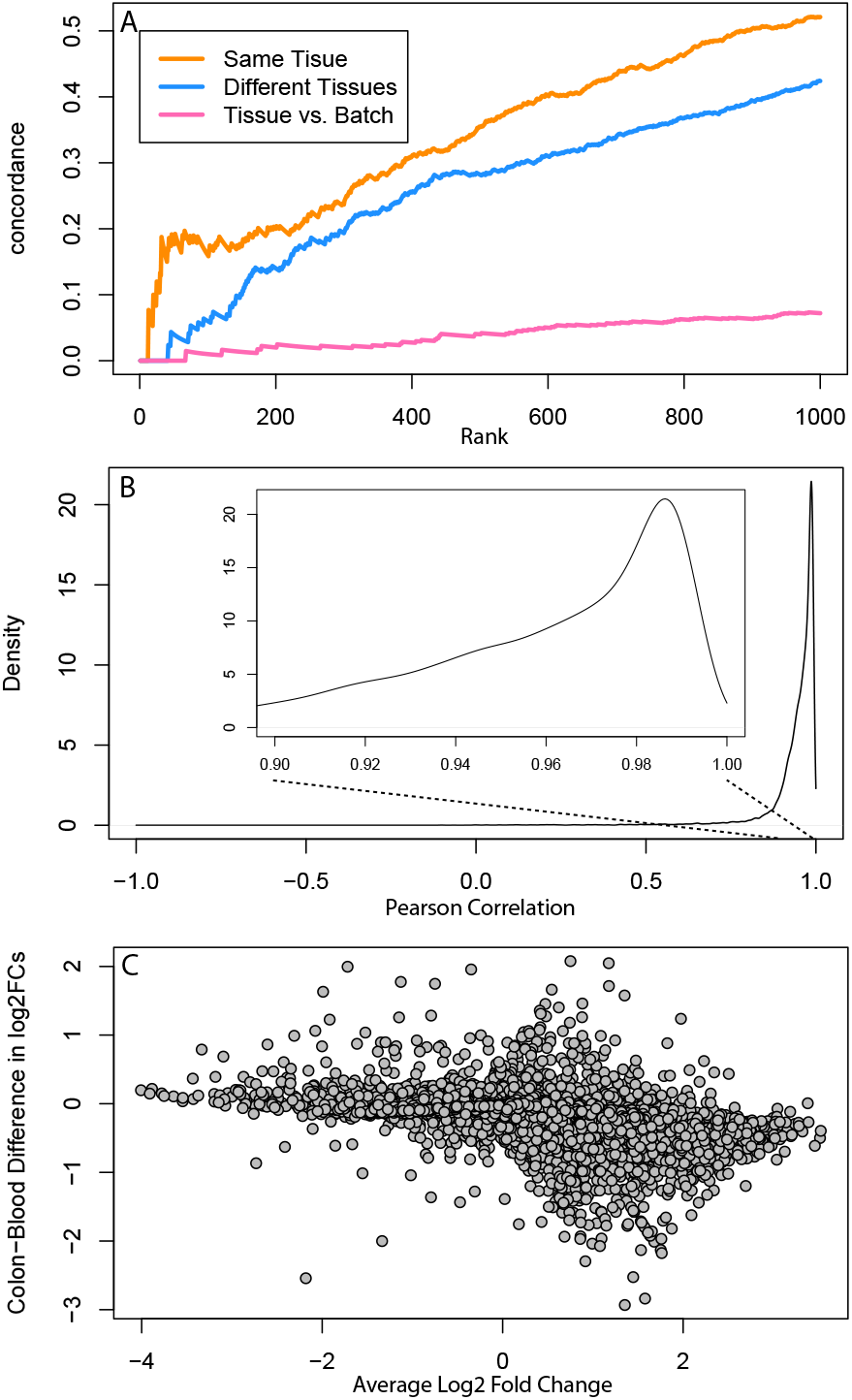
Meta-analysis and study comparison facilitated by recount A. A concordance at the top plot showing comparisons between a meta-analysis tissue comparison of whole blood and colorectal tissue in data from the sequence read archive and the GTEx project. When comparing the same tissues there is strong concordance between differential expression results on public data and GTEx, less when different tissues are compared, and almost none when comparing different analyses. **B.** The distribution of correlations between gene expression estimates for GTEx V6 from the GTEx portal and the counts calculated in recount. The gene expression counts are highly correlated between both quantifications. **C.** An MA-plot comparing the fold changes for differential expression between colon and whole blood using the quantifications from GTEx and from recount. Most genes have similar fold change between the two analyses.

### 2.3 Use case 2: GTEx comparison

One of the largest collections of RNA sequencing data currently available are data from the GTEx project consisting of 9,662 samples from over 250 individuals [19]. The recount collection includes the RNA-seq data from GTEx processed using the same pipeline as all other samples from SRA. The exon, gene, and junction counts are available from recount in the form of both tab-delimited files and analysis-ready Bioconductor objects.

We downloaded the official release of the gene counts from the GTEx portal (which were based on read counting) and compared them to our genes counts (which were based on base-level coverage). The gene expression levels we estimate using the recount pipeline have a median (IQR) correlation of 0.96 (0.93, 0.98) with the V6 release from GTEx (Figure 1B). We performed a differential expression analysis comparing colon and whole blood samples. Differential expression analysis using the gene expression measurements from recount match the results using the V6 release from the GTEx portal (*r*^2^ = .93 between fold changes for recount and GTEx v6 counts, Figure 1C). The advantage of using the recount version of the GTEx data is that they are already processed identically to tens of thousands of SRA samples and can be easily integrated to perform more comprehensive analyses as we have shown in previous examples.

### 2.4 Use case 3: Multi-level differential expression analyses

To demonstrate the ease with which differential gene expression analyses can be carried out in recount, we lastly performed differential expression (DE) analysis at the gene, exon, exon-exon-junction, and expressed region levels from data generated to determine the transcriptomic differences between breast cancer subtypes. In this first analysis, HER2-positive and triple negative breast cancer (TNBC) samples were selected from study SRP032789 (TNBC, n = 6; HER2-positive, n = 5) [20], and feature-level expression at genes, exons, junctions, and expressed regions were extracted (see Supplementary Methods). DE analysis at the gene-level identified 1,611 differentially expressed genes (q *<* 0.05) with 933 genes demonstrating decreased expression and 678 increased expression in TNBC relative to HER2-positive breast cancer. At the exon level, 23,647 exons demonstrated differential expression (q *<* 0.05; 11,218 downregulated and 12,429 upregulated). Finally, 19,805 exon-exon junctions were differentially expressed (q *<* 0.05, 18,073 downregulated and 1,732 upregulated). DE analysis identified 35,809 differentially expressed regions (q *<* 0.05, 17,170 downregulated and 18,639 upregulated). Of these significant DERs, 6,613 do not overlap any annotated exons, demonstrating that 18% of DERs detected would not be reported using annotation-dependent methods of expression estimation. Figure 2C highlights an example region in which differential expression occurs outside of any annotated protein-coding gene on Chromosome 3.

**Figure 2:**
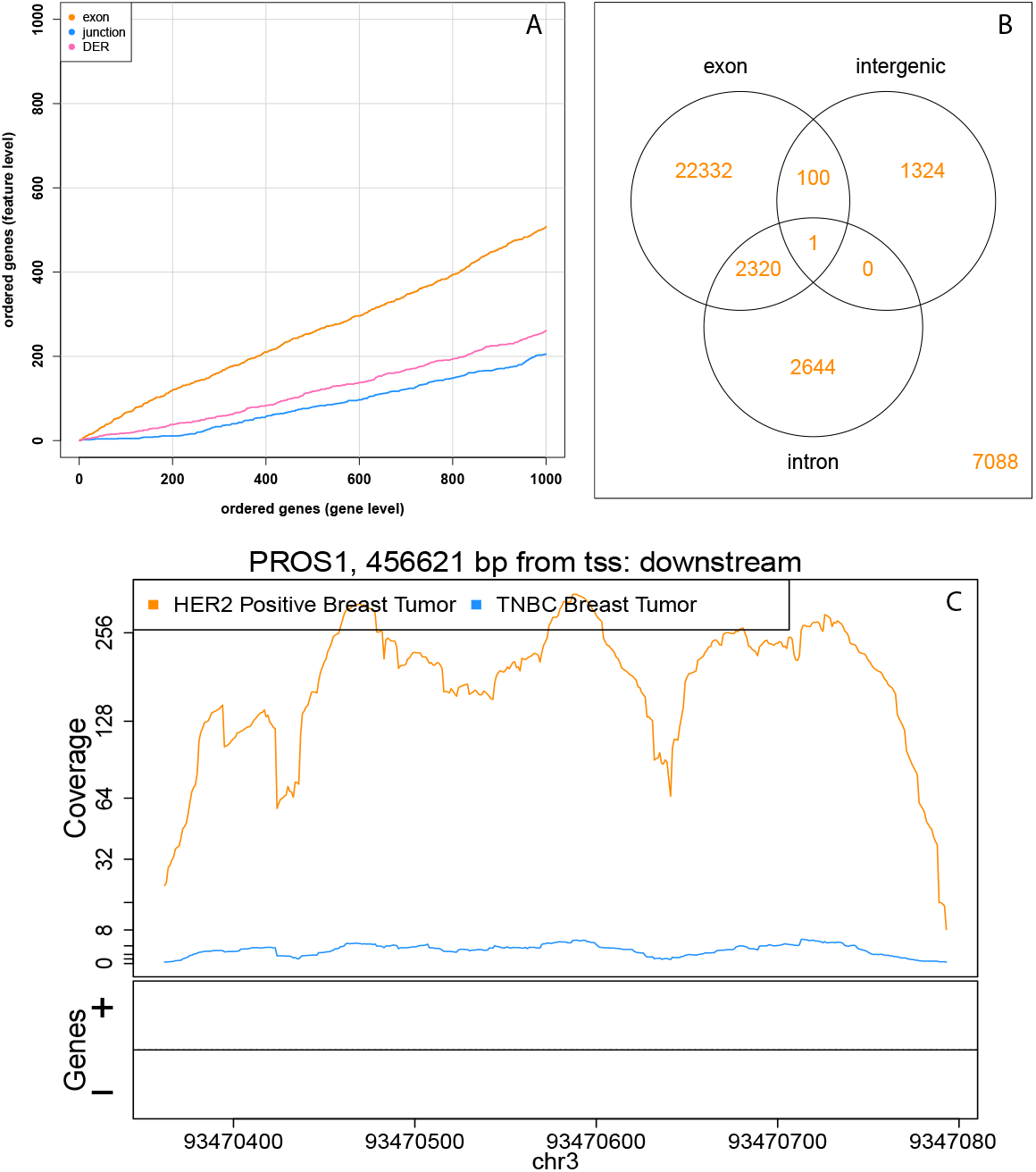
Multi-level differential expression analysis is facilitated by recount A. A concordance at the top plot showing concordance between a gene-level analysis, an exon-level analysis, a junction level analysis, and an expressed region level analysis (DER). All of these analyses are possible and comparable with recount. **B.** A Venn diagram showing the number of expressed regions detected that overlap exons, intergenic regions, and intronic regions, including expressed regions that overlap multiple annotation types. Differential expression occurs outside of previously annotated protein-coding regions. **C.** An example of a region showing differential expression between breast tumor subtypes where there is no annotated gene present. The lines show the average coverage in each group across samples.

We then summarized junctions and exons at the gene-level using the resulting DE p-values, and 67% of the top 100 genes were shared across the gene and exon-level analyses (Figure 2A). In comparison, expressed regions and exon-exon junction analyses only shared 18% and 5% of the top 100 genes, respectively. Furthermore, to validate the differential expression findings, we compared the gene level results from study SRP032789 to an independent study (SRP019936; TNBC, n = 8; HER2-positive, n = 7) [21] (see Supplementary Methods). DE analysis was carried out as described above, identifying 3,197 genes as differentially expressed (q *<* 0.05, 1,728 downregulated and 1,469 upregulated). Given the low concordance (8% among the top 1000 genes) between these results and those from study SRP032789, we then applied independent hypothesis weighting (IHW) [22] across the two studies, which slightly improved replication rates. While sample size is limited in these two studies and thus likely thwarts our ability to see a huge increase in power using IHW, example code for IHW in recount is provided for application to other data sets.

## 3 Discussion

By producing summaries at multiple levels of detail, recount enables a range of downstream analyses. The summaries are concise and easy for users to download and use. Achieving an appropriate balance between conciseness and queryability is an important design challenge for any effort that seeks to make public data more usable for researchers.

All `recount` summaries are produced with analysis pipelines that are both reproducible and annotation-agnostic. Gene annotations are used only to label summarized data post analysis, and not to align reads or discover splice junctions. Downstream analyses are therefore fully aware of unannotated splicing events.

Other efforts have been made to summarize public gene expression data. The Expression Atlas [23] provides final results queryable only at the gene level, Toil focuses on curated data [24], and other efforts have focused primarily on cancer [25].

`recount`, by contrast, covers a broad range of projects and produces summarized objects that can be further analyzed in a variety of ways - including directly with Bioconductor using the `recount` package.

The `recount` website is located at https://jhubiostatistics.shinyapps.io/recount/. The recount Bioconductor package is available under the Artistic 2.0 open source license and is available at https://github.com/leekgroup/recount.

## 4 Online Methods

### 4.1 Alignment

GTEx and public SRA samples were selected by searching the SRA website. Samples that could not be downloaded using fastq-dump were eliminated as discussed in Supplementary Methods.

Samples were aligned in a spliced fashion to the hg38 assembly of the human genome using Rail-RNA. Alignments were performed in batches on computer clusters rented from the Amazon Web Services Elastic MapReduce service. The alignment pipeline was divided into two phases, where the first phase (“preprocessing”) downloads and reformats the data and the second phase performs spliced alignment. Outputs of the pipeline include, for each sample, a junction coverage file (similar to a TopHat “junctions.bed” file) and a BigWig file [26] containing a genomewide coverage vector. Further details are presented in the Rail-RNA study [27] and Supplementary Methods.

### 4.2 Tabulation

Gene and exon counts were compiled using the BigWig files output by Rail-RNA and the UCSC knownGene annotation. For exon counts, we first obtained a set of non-overlapping “unioned” exons. Gene and exon counts were compiled into per-project tables and RangedSummarizedExperiment objects. We expanded the tables with several metadata columns containing, for example, read count, paired-end status, GEO accession, and the tissue type as predicted by the SHARQ beta resource (http://www.cs.cmu.edu/∼ckingsf/sharq/).

### 4.3 Use cases

All R code used for the analyses performing the use cases is available from the website: http://leekgroup.github.io/recount-analyses/.

- Use case 1: Meta-analysis http://leekgroup.github.io/recount-analyses/example_meta/meta_analysis.pdf
- Use case 2: GTEx comparison http://leekgroup.github.io/recount-analyses/example_meta/compare_with_GTEx_reproducible.pdf
- Use case 3: Multi-level differential expression analyses Gene/exon: http://leekgroup. github.io/recount-analyses/example_de/recount_SRP032789.pdf and annotation agnostic: http://leekgroup.github.io/recount-analyses/example_de/recount_DER_SRP032789.pdf and validation of results in a second study http://leekgroup.github.io/recount-analyses/example_de/recount_SRP019936.pdf

## 5 Acknowledgments

We thank Carl Kingsford and Darya Filippova for their assistance in adding SHARQ metadata to recount. recount data is hosted on SciServer, a collaborative research environment for large-scale data-driven science. It is being developed at, and administered by, the Institute for Data Intensive Engineering and Science at Johns Hopkins University. SciServer is funded by the National Science Foundation Award ACI-1261715. For more information about SciServer, visit http://www. sciserver.org.

The Genotype-Tissue Expression (GTEx) Project was supported by the Common Fund of the Office of the Director of the National Institutes of Health. Additional funds were provided by the NCI, NHGRI, NHLBI, NIDA, NIMH, and NINDS. Donors were enrolled at Biospecimen Source Sites funded by NCI/SAIC-Frederick, Inc. (SAIC-F) subcontracts to the National Disease Research Interchange (10XS170), Roswell Park Cancer Institute (10XS171), and Science Care, Inc. (X10S172). The Laboratory, Data Analysis, and Coordinating Center (LDACC) was funded through a contract (HHSN268201000029C) to The Broad Institute, Inc. Biorepository operations were funded through an SAIC-F subcontract to Van Andel Institute (10ST1035). Additional data repository and project management were provided by SAIC-F (HHSN261200800001E). The Brain Bank was supported by a supplements to University of Miami grants DA006227 & DA033684 and to contract N01MH000028. Statistical Methods development grants were made to the University of Geneva (MH090941 & MH101814), the University of Chicago (MH090951, MH090937, MH101820, MH101825), the University of North Carolina - Chapel Hill (MH090936 & MH101819), Harvard University (MH090948), Stanford University (MH101782), Washington University St Louis (MH101810), and the University of Pennsylvania (MH101822). The data used for the analyses described in this manuscript were obtained from: the GTEx Portal on 11/21/15 and/or dbGaP accession number phs000424.v6.p1 on 11/30/15 - 12/04/15.

## Supplementary Methods and Results

### 1 Supplementary Results

### 2 Supplementary Methods

#### Selecting GTEx samples

On November 21, 2015, we queried the SRA website at http://www.ncbi.nlm.nih.gov/sra for RNA-seq samples in the GTEx project (accession: SRP012682). Precise search terms were (SRP012682) AND “strategy rna seq"[Properties]. A screenshot is available at https://github.com/nellore/runs/raw/master/gtex/SRA_GTEx_search_screenshot_ 6.37.16_PM_ET_11.21.2015.png. We used the “send to file” function to download search results to a table, available at https://github.com/nellore/runs/raw/master/gtex/SraRunInfo.csv. The table lists 9,795 run accessions, some of which were mmPCR-seq. We restricted our attention to the 9,662 mRNA-seq samples in the V6 release of the GTEx consortium.

#### Selecting SRA samples

Public SRA samples were selected from SRA as follows. On February 3, 2016, we queried the SRA website at http://www.ncbi.nlm.nih.gov/sra for publicly available human RNA-seq samples. Precise search terms were “platform illumina"[Properties] AND “strategy rna seq"[Properties] AND “human"[Organism] AND “cluster public"[Properties] AND “biomol rna"[Properties]. A screenshot is available at https://github.com/nellore/ runs/raw/master/sra/v2/hg38/SRA_RNA-seq_search_screenshot_3.33.19_PM_ET_02.03.2016. png. We used the “send to file” function to download the search results to a table, available at https://github.com/nellore/runs/raw/master/sra/v2/hg38/SraRunInfo.csv. The table lists 50,186 run accessions, each with metadata fields including layout (SINGLE or PAIRED) and number of reads (i.e., read pairs for paired-end samples).

#### Alignment with Rail-RNA

We used Rail-RNA v0.2.3 [1] to align all selected samples we could download to the *hg38* assembly. Alignment was performed using the Amazon Web Services Elastic MapReduce commercial cloud computing service. that is, standardized units of computing capacity.

Spot instances allow users to bid for excess computing capacity. If the fluctuating market price drops below a users’s bid, the instances could be lost, halting the computation. So saving money by bidding for spot instances comes with risk, and rather than aligning all samples in one batch, we distributed this risk by dividing alignment up [into batches]. For GTEx, we randomly divided the set of 9,662 samples up into 30 batches, each with 322 or 323 samples; for other SRA samples, we randomly divided the set of samples into 100 batches, each with 501 or 502 samples. Analysis of each batch was itself divided into (1) a preprocessing job flow, which uses SRA Tools fastq-dump (https://github.com/ncbi/sra-tools) to download compressed reads from the National Center for Biotechnology Information server and preprocess them, writing to Amazon’s cloud storage service S3; and (2) an alignment job flow. Every preprocessing job flow was run on a cluster of 21 m3.xlarge instances, each with 4 Intel Xeon E5-2680 v2 (Ivy Bridge) processing cores and 15 GB of RAM. Every alignment job flow was run on a cluster of 81 c3.8xlarge instances, each with 32 Intel Xeon E5-2680 v2 (Ivy Bridge) processing cores and 60 GB of RAM. Our GTEx alignment runs may be reproduced by following instructions at https://github.com/nellore/runs/blob/ master/gtex/README.md. Our alignment runs may be reproduced by following instructions at https://github.com/nellore/runs/blob/master/sra/v2/README.md. Alignment of GTEx data is described in more detail in the supplementary material of [2].

#### Incomplete and invalid input data

For the analysis of the public SRA samples, some samples could not be downloaded, due to failures of the fastq-dump software. The issue persisted even when we attempted to restart the fastq-dump process. Samples exhibiting this issue can be excluded from analysis. These samples are listed here: https://github.com/nellore/runs/raw/master/ sra/v2/hg38/NOTES. That file also describes two other preprocessing issues that led us to exclude a few other samples from analysis: (1) sequence input encoded in a manner we did not recognize; (2) miscellaneous errors reported by fastq-dump.

In addition, for each of a small number of both GTEx and public SRA samples, fastq-dump would return success (an exit code of 0) when an error had occurred and the number of reads output by the tool disagreed with the number of reads from SRA metadata given in the SraRunInfo.csv file for that sample. So for each sample, we compared the number of reads ingested by Rail-RNA with the number of reads reported in SraRunInfo.csv. Sometimes, the sample was listed as paired-end when the number of mates (i.e., twice the number of read pairs listed in SRARunInfo.csv) was exactly twice the number of reads Rail-RNA processed; other times, the sample was listed as single-end but the number of reads Rail-RNA processed was greater than the number of reads listed. In these cases, we made note of our suspicion that the library layout listed on SRA was incorrect.

The table https://github.com/nellore/runs/raw/master/gtex/incomplete.tsv (respectively, https://github.com/nellore/runs/raw/master/sra/v2/incomplete.tsv) lists all GTEx (respectively, public SRA) samples that either could not be downloaded or were incompletely downloaded, and it includes these suspected layout misreports in a column. This table was generated by the script https://github.com/nellore/runs/raw/master/gtex/incomplete.py (re-spectively, https://github.com/nellore/runs/raw/master/sra/v2/incomplete.py) for GTEx (respectively, public SRA) samples.

#### Counting genes and exons

When creating the count tables, genes and exons (features) were determined using TxDb.Hsapiens.UCSC.hg38.knownGene package version 3.1.3 [3]. For each gene, we first obtained a set of non-overlapping exonic intervals by taking the “union” of the gene’s annotated exons. We call these “unioned exons”. We summed per-base coverage across each unioned exon using bwtool version 1.0 [4]. We obtained gene-level counts by summing per-base coverage totals of its constituent unioned exons. For each SRA project and each feature type we created a RangedSummarizedExperiment object [5] containing columns for:

- SRA study id
- SRA sample id
- SRA experiment id
- SRA run id
- read counts as reported by SRA
- number of reads downloaded from SRA and subsequently aligned with Rail-RNA
- proportion of reads reported by SRA that aligned
- whether the sample was paired-end or not
- whether SRA likely misreported the paired-end label (as explained above)
- mapped read count by Rail-RNA
- coverage area under the curve (AUC), i.e. total number of aligned bases not including soft-clipped bases
- tissue type as predicted by SHARQ (http://www.cs.cmu.edu/∼ckingsf/sharq/)
- cell type as predicted by SHARQ
- biosample submission date
- biosample publication date
- biosample update date
- average read length as reported by SRA
- GEO accession id
- sample title as reported by GEO
- sample characteristics as reported by GEO
- name of the coverage BigWig file

For each sample we normalized the coverage to 40 million 100 base-pair reads using the AUC information. We then computed the base-level coverage sum for each project using the sample normalized coverage with bwtool version 1.0 [4] and divided the coverage sum by the number of samples in the project resulting in the project mean coverage. We then stored the mean coverage for each SRA project in a BigWig file [6]. The mean coverage can then be used to identify expressed regions with derfinder [7] via the recount R package. The code and log files for creating these files as well as the recount website are available at https://github.com/leekgroup/recount-website. **Exon-exon junction output processing.** We postprocessed Rail-RNA’s cross-sample junction TSVs to obtain two gzip-compressed files for each project on SRA: [SRA project accession number].junction id with transcripts.bed.gz, a file in BED format that contains junction coordinates, assigns each junction a unique identifier in the name column, and also lists which transcripts in GENCODE v24 [8] contain the junction or its corresponding donor and acceptor site; and [SRA project accession].junction coverage.tsv.gz, which for each junction identifer gives the number of reads that map across the junction in each sample in the project. This postprocessing was performed by two scripts run in succession: https://github.com/nellore/runs/blob/master/sra/v2/hg38/junctions_by_project.py and https://github.com/nellore/runs/blob/master/sra/v2/hg38/add_knowngene.py.

For each project, we created a RangedSummarizedExperiment using the sample metadata as in the exon and gene cases while annotating the exon-exon junctions with the:

- exon-exon junction id
- GENCODE v24 transcript id
- transcript name, gene id and gene symbol based on TxDb.Hsapiens.UCSC.hg38.knownGene
- class: annotated, exon skip, alternative end, fusion, novel
- proposed gene id and gene symbol based on TxDb.Hsapiens.UCSC.hg38.knownGene.

The code is available at https://github.com/leekgroup/recount-website.

#### Breast Cancer Use Case

Principal component analysis (PCA) was run to identify any sample outliers. All samples were retained for downstream analyses. The 17,874 genes with an average normalized count greater than five across samples were included for downstream analysis. Similarly, 204,559 exons and 45,933 exon-exon junctions with reads overlapping those genes were also included for DE analysis. Finally using this data set, we further included analysis of data at the level of expressed regions to highlight the utility of summarizing expression in an annotation-agnostic manner. At the expressed region level, 291,247 regions with an average normalized read count greater than five across samples were included for DE analysis. DE between the two cancer subtypes was carried out using limma and voom [9]. For multiple comparison correction we calculated q-values from the observed p-values after estimating the proportion of differentially expressed genes, exons, or exon-exon junctions in the experiment [10]. Features with calculated q-values smaller than 0.05 between different tissue types were declared statistically significant. To compare the results across these levels of data, we used Simes’ rule [11] to calculate a gene-based p-value for all exons, exon-exon junctions, and differentially expressed regions that overlap genes. We then computed rank-based concordance between the gene and either the exon, junction, or expressed region level results.

In the replication dataset, count data were filtered as before and PCA was utilized to identify global sample outliers. One TNBC tumor sample was identified as an outlier and removed from analysis. IHW uses a set of independent weights for each gene in one study to weight the hypothesis tests in the second study to increase power to detect differences. Here, for each gene the absolute value of the test statistic from study SRP032789 were used as weights for DE analysis in study SRP019936. This is akin to using an empirical prior and treating the second study as a validation of the first.

